# Cerebrovascular reactivity increases across development in multiple networks as revealed by a breath-holding task: a longitudinal fMRI study

**DOI:** 10.1101/2023.01.05.522905

**Authors:** Donna Y. Chen, Xin Di, Bharat Biswal

## Abstract

Functional magnetic resonance imaging (fMRI) has been widely used to understand the neurodevelopmental changes that occur in cognition and behavior across childhood. The blood-oxygen-level-dependent (BOLD) signal obtained from fMRI is understood to be comprised of both neuronal and vascular information. However, it is unclear whether the vascular response is altered across age in studies investigating development in children. Since the breath-hold task is commonly used to understand cerebrovascular reactivity in fMRI studies, it can be used to account for developmental differences in vascular response. This study examines how the cerebrovascular response changes over age in a longitudinal children’s breath-hold dataset from the Nathan Kline Institute (NKI) Rockland Sample (ages 6 to 18 years old at enrollment). A general linear model (GLM) approach was applied to derive cerebrovascular reactivity from breath-hold data. To model both the longitudinal and cross-sectional effects of age on breath-hold response, we used mixed effects modeling with the following terms: linear, quadratic, logarithmic, and quadratic-logarithmic, to find the best-fitting model. We observed increased breath-hold BOLD signal in multiple networks across age, in which linear and logarithmic mixed effects models provided the best fit with the lowest Akaike Information Criterion (AIC) scores. This shows that the cerebrovascular response increases across development in a brain network-specific manner. Therefore, fMRI studies investigating the developmental period should account for cerebrovascular changes which occur with age.

## 1. Introduction

The developmental period is a critical time when distinct brain regions undergo rapid growth. With advancements in functional magnetic resonance imaging (fMRI) equipment and analytic methods, we have a better understanding of the developing brain (Davidson et al., 2003; Van Horn and Pelphrey, 2015). fMRI studies have shown differences in brain organization between children and adults, with children having a more localized brain network structure and adults having a more segregated network structure (Fair et al., 2009). Additionally, fMRI studies have revealed developmental changes in cognitive control, such as inhibition response and working memory (Luna et al., 2010), changes in emotional processing in adolescence (Somerville et al., 2010), and altered brain connectivity patterns in children with neurodevelopmental disorders such as autism spectrum disorder (Uddin et al., 2013).

The blood-oxygen-level-dependent (BOLD) fMRI signal is an indirect measure of neuronal activity and is based on the theory of neurovascular coupling, in which an increase in neuronal activity is followed by an increase in blood flow to the particular brain region of activation (Iadecola, 2017). Age-related disruptions in neurovascular coupling have been observed (Abdelkarim et al., 2019; Galiano et al., 2020; Tsvetanov et al., 2015; West et al., 2019) and may be due to a range of potential mechanisms such as an increase in inflammatory signaling, oxidative stress, vascular remodeling due to hypertension, cerebral amyloid angiopathy, arterial stiffening, or the breakdown of the blood brain barrier, which disrupt vasomotor activity (Zimmerman et al., 2021). Furthermore, decreased vascular reactivity in older populations influences the BOLD fMRI signal and may be a confounding factor in studies investigating aging (Kannurpatti et al., 2014). This is important to consider in age-related fMRI studies since differences observed may be mediated by vascular changes rather than neuronal changes. However, little is yet known about the neuronal and vascular contributions of the BOLD signal during childhood. The brain’s hemodynamic response is faster in children than that of adults, since the blood vessel walls tend to stiffen with age, along with changes that occur in the heart rate and respiratory rate which are faster in children (Thomason et al., 2005). Church and colleagues suggest that the developmental differences observed in fMRI studies are not vascular, since the percent BOLD signal change peaks were different among different tasks (Church et al., 2010). However, Schmithorst and colleagues analyzed the ratio of BOLD signal to relative cerebral blood flow (CBF) changes and found increased neurovascular coupling in the middle temporal gyri and the left inferior frontal gyrus with age in children aged 5-18 years (Schmithorst et al., 2015). This supports that the BOLD signal changes may not be driven primarily by neuronal changes, but rather a strong vascular component that influences the changes seen in developmental fMRI studies. Therefore, the vascular properties of the brain contribute to the fMRI signal and need to be further investigated as they may act as a confound.

In order to study the vascular properties of the brain with fMRI, the breath-holding task can be used since it focuses more on the vascular contribution of the BOLD signal rather than the neuronal feature. There are many neurovascular processes that occur in the brain which contribute to the BOLD signal, such as glial cell interactions with the vessel wall (Logothetis and Pfeuffer, 2004), the control of capillary walls by pericytes (Mark et al., 2015), and signaling between neurons and astrocytes to control CBF (Drew, 2019). However, BOLD signal changes under breath-holding are the result of dilation due to increased arterial CO2 concentration rather than being primarily neurotransmitter-mediated (Chen and Pike, 2009), and thus provides a measure of vascular function. It is important to note that the BOLD signal amplitude is also modulated by baseline states of cerebral blood volume (CBV), cerebral metabolic rate of oxygen (CMRO2), and cerebral blood flow (CBF), therefore, the breath-hold response may be modulated by these factors as well and not only the degree to which vessels dilate. The BOLD signal depends on baseline concentrations of deoxy-hemoglobin and total oxy-hemoglobin, which are mediated through CBF and CBV, respectively (Mark et al., 2015). Furthermore, hypercapnia induced changes in the BOLD signal are often expressed in terms of percentage signal changes from baseline BOLD signal, thus being dependent on the baseline parameters (Bhogal et al., 2016). During the breath-holding task, participants are commonly asked to perform a block-design experiment consisting of blocks of breath-holding ranging from 15-20s, interleaved with blocks of normal-paced breathing ranging from 15-30s (Kastrup et al., 1999b; Kwong et al., 1995; Stillman et al., 1995). This allows researchers to assess the cerebrovascular response to a vasoactive stimulus, such as increased carbon dioxide levels. As one holds their breath, there is a build-up of arterial carbon dioxide concentration and a decrease in pH levels, which leads to the vasodilation of blood vessels that occurs globally in the brain (Bright et al., 2011; Kety and Schmidt, 1948). Other methods used to study the vascular contribution of the BOLD signal may be more involved, such as the intake of acetazolamide or the controlled inhalation of different levels of carbon dioxide gas (Bright et al., 2009; Bruhn et al., 1994; Mukherjee et al., 2005).

Thomason and colleagues reported lower BOLD fMRI signal to noise ratio (SNR) in children compared to that of adults during the performance of a breath-hold task and found a reduced volume of response in children (Thomason et al., 2005). They also found a heterogeneous vascular response pattern across the brain, which was similar between children and adults. This shows that the BOLD fMRI signal differences reported between groups in task-based studies may not be reflective of differences in neuronal activity but rather reflects the differences in vascular response between groups. However, a limitation of the study includes its cross-sectional design in which comparisons were made between two groups: children (7 to 12 years) and young adults (18 to 29 years). The effect of age on vascular activity may not necessarily be linear but rather have complex trajectories (Faghiri et al., 2019; Gracia-Tabuenca et al., 2021). Therefore, investigating age in one group containing a range of ages may be preferred over comparing results between distinct groups. Furthermore, developmental changes to the BOLD hemodynamic response function (HRF) have been shown in a study comparing neonates to adults, in which neonates had a lower HRF positive peak and longer time-to-peak duration than adults in response to a somatosensory stimulus (Arichi et al., 2012). Additionally, in a study investigating children with epilepsy, significantly longer peak times of the HRF was found in children aged 0 to 2 years old; however, no differences in the HRF were observed between older children and adults with epilepsy (Jacobs et al., 2008). It is not yet clear how the cerebrovascular reactivity (CVR) pattern changes across development.

Leung and colleagues characterized cerebrovascular reactivity from childhood to early adulthood (9-30 years, N=34), and found an increase in CVR peaking in the mid-teens, followed by a subsequent decrease into adulthood (Leung et al., 2016). This shows the piece-wise trajectory of CVR from childhood to early adulthood; however, the study did not investigate potential regional differences in CVR across the ages and the study implemented a cross-sectional design rather than a longitudinal design. Urback and colleagues found the largest amplitude in CVR in the frontal and visual regions in healthy adolescents (13-19 years), which was significantly higher than the CVR in the parietal and temporal regions (Urback et al., 2018). Thomason and colleagues also show in a group of children from 7 to 12 years of age, that the percent fMRI signal change during breath-holding was highest in the visual and motor regions of the brain (Thomason et al., 2005). There have been studies that investigated regional CVR differences in adults, which reported a similar pattern to that of children, with the exception of CVR in the frontal region being the highest in children (Kastrup et al., 1999a). Since the frontal regions are one of the last to mature (Giedd, 2008), we hypothesize that CVR would increase in this region until late adolescence, reflective of delayed cerebrovascular development (Urback et al., 2018). Therefore, we expect the developmental trajectories of CVR to be dependent on the specific brain region under study.

Non-linear developmental trajectories from childhood to early adulthood have been reported in both structural and functional brain maturation (Dosenbach et al., 2010; Faghiri et al., 2019; Giedd et al., 1999; Lebel and Beaulieu, 2011; Tamnes et al., 2017; Wierenga et al., 2014); however, the developmental trajectories of CVR in children is poorly understood. Leung and colleagues showed a piece-wise linear trend in CVR across children, in which CVR increased after 9 years of age, peaked at around 14.7 years, and then subsequently decreased to 30 years of age (Leung et al., 2016). This study was one of the few studies which investigated CVR in children and suggests a complex nonlinear developmental trajectory in CVR in children. There have been other studies which report nonlinearities in brain development, particularly in that of structural brain development. Cortical gray matter volume has been shown to increase in early childhood, peaking at different ages depending on the brain region of interest (Wierenga et al., 2014), and then typically decreases in adolescence (Giedd et al., 1999). The higher-order association cortices mature at a delayed rate than the somatosensory and visual cortices, revealing not only a nonlinear process, but also different region-specific trajectories (Gogtay et al., 2004). Shaw and colleagues found peak maturation and developmental trajectories to differ based on different areas of the brain, such that the superior frontal gyri followed a cubic curve, the insula a quadratic curve, while the orbitofrontal cortex followed a linear curve (Shaw et al., 2008). Furthermore, nonlinear developmental trajectories have also been observed in functional fMRI studies. Faghiri and colleagues found a U-shaped pattern of connectivity across early childhood through adolescence, speculating that in mid-adolescence, connectivity weakens (Faghiri et al., 2019). However, despite the strong evidence of nonlinear trajectories observed in neurodevelopment across children, it is not clear whether CVR also follows similar trends in development, as there is a lack of studies investigating the developmental trajectories of CVR (Ainslie and McManus, 2016; Ellis and Fluck, 2016).

The goal of the current study was to investigate the breath-hold response in children across different ages of development ranging from 6-20 years, using a longitudinal fMRI dataset from the Nathan Kline Institute (NKI) (Tobe et al., 2022). This dataset includes both longitudinal and cross-sectional data, which allows us to better understand within-subject and between-subject developmental trajectories, respectively. Here, we investigate cerebrovascular reactivity changes across development by quantifying the response to carbon dioxide in multiple brain regions during breath-holding. We hypothesize that cerebrovascular reactivity will show the greatest changes in the frontal and visual regions of the brain and follows a nonlinear trend in children, in which there is an initial increase in cerebrovascular reactivity, followed by a plateau as cerebrovasculature reaches maturity.

## 2. Materials and Methods

### 2.1. Dataset

The Nathan Kline Institute Rockland Sample (NKI-RS) dataset is a community-ascertained collection of multimodal MRI, cognitive, behavioral, and phenotypic data across the lifespan (https://data.rocklandsample.rfmh.org/). For this study, we focused on the NKI-RS Longitudinal Discovery of Brain Development Trajectories sub-study (N=369) (Tobe et al., 2022), which implements a longitudinal design to study connectome development in children. The age at enrollment ranged from 6 to 17 years and the oldest age at completion of the studies is 20 years. We analyzed the breath-hold fMRI data for all healthy subjects, to study vascular activity in the brain due to a vasoactive hypercapnic stimulus. The healthy control (HC) group was defined as having no diagnosis or condition on Axis I of the Diagnostic and Statistical Manual of Mental Disorders (DSM-IV) (Bell, 1994).

For the breath-holding paradigm there were two different task designs implemented: one with an 18s breath-hold duration and the other with a 15s breath-hold duration. There were a total of 7 blocks of breath-holding each lasting either 18 or 15 seconds, interleaved with 18 second blocks of self-paced breathing. During breath-holding, the participants were shown a circle on a computer screen of decreasing size indicating the duration of breath-holding left. Since this was a longitudinal study, three timepoints were collected from some of the participants: baseline (BAS1), first follow-up (FLU1), and second follow-up (FLU2) (Figure 1). The follow-up sessions were spaced either 12 months or 15 months apart. Some participants enrolled in the 0/12/24 month track while others enrolled in the 0/15/30 month track. We aggregated the subjects on the two tracks together and included subjects who performed either 15s or 18s breath-hold paradigms, to maximize the number of subjects to be analyzed. However, only subjects who had consistency in the breath-hold paradigm across different sessions were included in this study, since longitudinal modeling was implemented. For those who had inconsistencies in the type of breath-hold paradigm performed (either 15s or 18s) from one session’s scan to the next, only the last session’s data was kept for that particular subject, therefore, that subject would only have one timepoint of data. The NKI study included participants based on the following criteria: 1) ages 6 to 20.5 years (ages 6 to17.9 years at enrollment), 2) children who have the capacity to understand and provide informed consent, 3) children aged 6-17.9 years who can sign assent and parent/guardian who can sign informed consent, 4) fluent in English, and 5) proof of residency in Rockland County, Orange County, Westchester County, NY, or Bergen County, NJ (Tobe et al., 2022).

**Figure 1.**
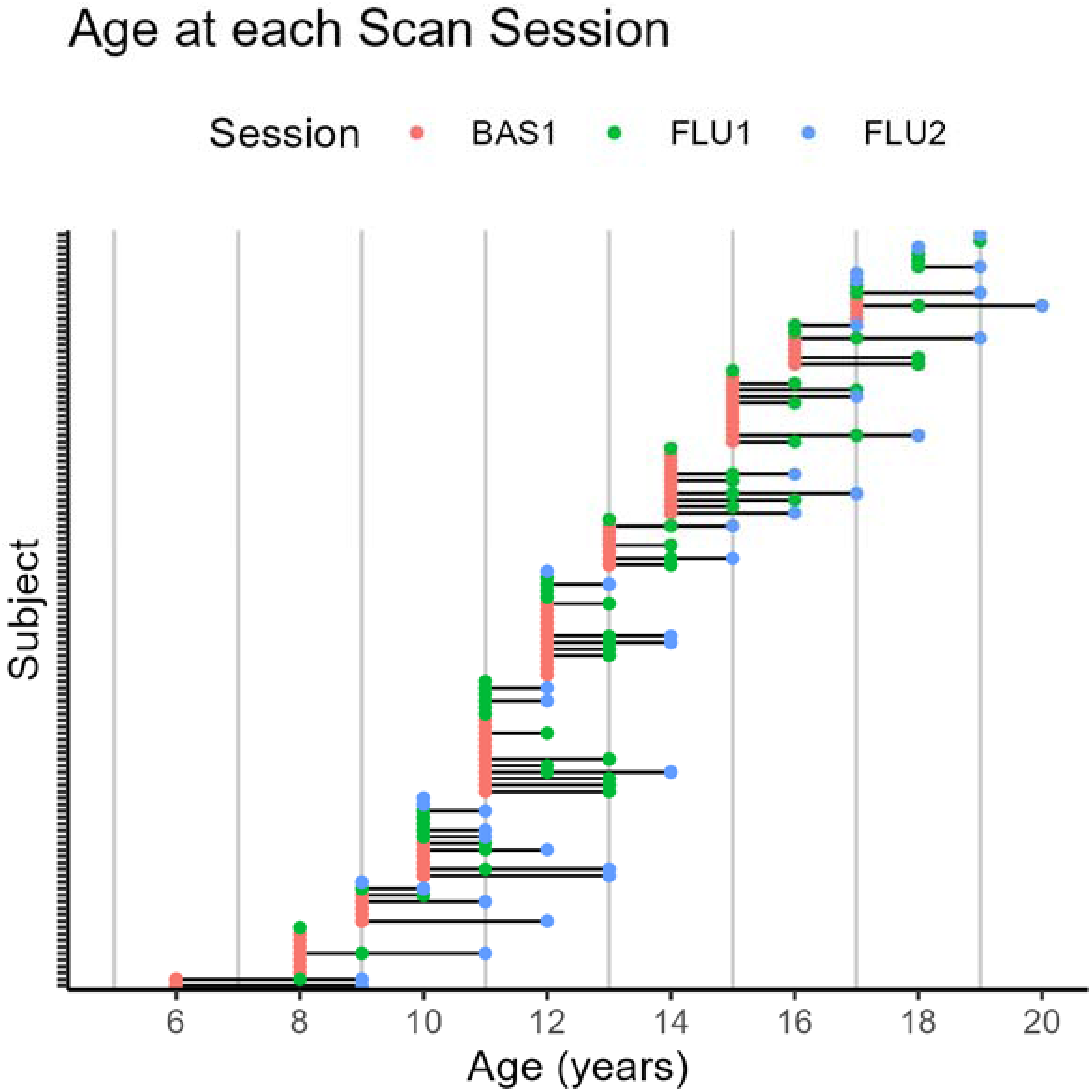
Age distribution at each scan session. Baseline (BAS1) sessions are denoted in red color, first follow-up (FLU1) in green, and second follow-up (FLU2) in blue. The black horizontal lines connecting the dots represent a single subject who completed multiple scan sessions.

### 2.2. Imaging Parameters

All MRI scans were obtained using a 3.0T Siemens TIM Trio at the Nathan Kline Institute, in which a 32-channel head coil was used. The acquisition time for the T1-weighted image was 4 minutes and 18s, with 176 slices, a TR of 1900ms and TE of 2.52ms. The acquisition time for the breath-hold fMRI data was 4 minutes and 30 seconds, with 64 slices, 2×2×2 mm^3^ resolution, a field of view (FOV) of 224 mm, % FOV phase at 100%, matrix size 112×112, a flip angle of 65 degrees, multiband acceleration factor of 4, TR of 1400ms, and TE of 30ms (Tobe et al., 2022).

### 2.3. Preprocessing

The data were preprocessed using Statistical Parametric Mapping (SPM12) (Ashburner et al., 2014) in MATLAB version 2021a (Mathworks, Natick, MA). First, each participants’ anatomical MRI image was segmented into gray matter, white matter, and cerebrospinal fluid. Each participants’ functional images were realigned to the mean functional image. For all participants, the framewise displacement values of translation and rotation were calculated to determine which participants had excessive head-motion. Any participants with a mean translation or mean rotation of greater than 0.3 mm were excluded from the study. Similarly, any participant with a maximum translation or maximum rotation greater than 2 mm was excluded from the study (Figure 2). These thresholds were defined by visualizing the framewise displacement values for all subjects, while taking into consideration the fMRI isotropic voxel size of 2 mm. Each subject’s realigned functional images were coregistered to their skull-stripped anatomical image. Subsequently, normalization was performed using SPM12’s diffeomorphic anatomic registration through an exponentiated lie algebra algorithm (DARTEL) (Ashburner, 2007). A study-specific template for DARTEL was first generated using each subject’s segmented anatomical images. Then, DARTEL normalization was performed to warp each subject’s coregistered functional images to the study-specific template. Smoothing was also performed using a Gaussian kernel with a full-width-half-maximum (FWHM) of 6 x 6 x 6 mm^3^. Lastly, all fMRI scans were visually inspected for poor data quality such as artifacts or missing brain regions on the image.

**Figure 2.**
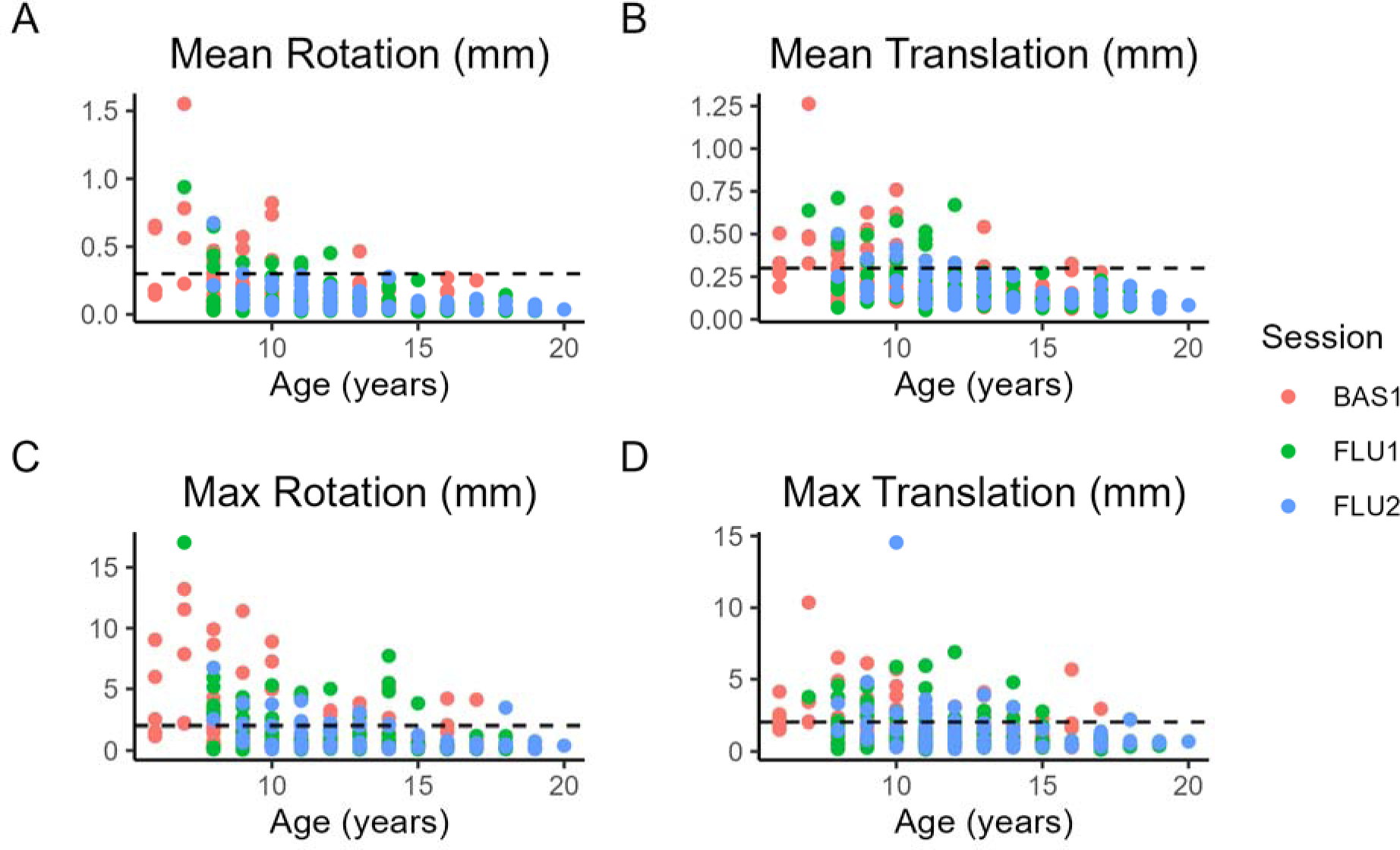
Framewise displacement values calculated for all participants for each age. The following framewise displacement values were calculated: A) mean rotation, B) mean translation, C) max rotation, and D) max translation. For mean values, a cut-off of 0.3 mm was chosen whereas for maximum values, a cut-off of 2 mm was chosen, as shown by the dashed lines.

### 2.4. Independent Component Analysis

Independent component analysis (ICA) was performed on a group-level for all breath-holding sessions together and then back-reconstructed on a subject-level to obtain subject-level ICA network maps. This was performed using the Group ICA of fMRI Toolbox (GIFT) (Rachakonda et al., 2007). Using the GIFT toolbox, the Infomax algorithm was used with default parameters applied, and the images were then z-scored with a threshold applied at z = 1.96. We obtained 25 independent components, and 12 of the 25 components were visually identified as meaningful brain networks rather than noise, thus were kept for further analyses. By using ICA, we were able to obtain spatially distinct networks, and thus reduced the number of brain areas of interest. It is important to note that these networks were defined using breath-holding data, rather than task or resting-state data, thus are not “functional brain networks”, but represents a more physiological or vascular network. Chen and colleagues show that physiological networks appear to mimic neuronal networks, and physiological signals such as respiratory volume and heart rate may give rise to the correlated signals across different brain regions as seen in large-scale resting-state brain networks (Chen et al., 2020). These physiological networks may provide meaningful characterizations of vascular anatomy. A network-level approach rather than a voxel-wise approach allows us to reduce the dimensionality of the data and reduce issues with multiple testing. Additionally, white-matter regions were not of interest in our analyses, since gray matter has up to 3 times greater CVR than white matter (Rostrup et al., 2000; Thomas et al., 2014) and the ratio of gray matter to white matter is higher in children than in adults (Marcar et al., 2004).

### 2.5. General Linear Model Analysis

To obtain a measure of cerebrovascular reactivity, a general linear model (GLM) was applied to the back projected ICA mean time-series data from all subjects. GLM was performed in MATLAB v2021a using SPM12’s canonical hemodynamic response function convolved with the task design. Prior to GLM, the data were standardized into z-scores (Rachakonda et al., 2007) for each network and each subject, in which the mean of the network’s mean time-series data was subtracted from the value at each timepoint and then this value was divided by the standard deviation of the mean time-series, resulting in a mean of 0 and standard deviation of 1 for the z-scored mean time-series data. GLM was performed for each of the 12 networks. The beta weight associated with the breath-hold task represents the breath-hold task response (Bright and Murphy, 2013). We performed GLM since we had two different breath-hold (BH) task designs – one with an 18s breath-holding period and another with a 15s breath-holding period. In the GLM, the design matrix was different between the 18s and 15s groups, to account for the different task design paradigms. The 18s breath-hold paradigm had a breath-hold stimulus onset at the following times: 18, 54, 90, 126, 162, 198, and 234 seconds, whereas the 15s breath-hold paradigm had a breath-hold stimulus onset at the following times: 18, 51, 84, 117, 150, 183, and 216 . A delay of 16s was added to both task designs to account for the delay associated with the breath-hold response.

### 2.6. Mixed Effects Model

Developmental effects of breath-holding activity were analyzed using a mixed-effects modeling approach. In our mixed effects model, the “age” variable was treated as a fixed effect, while the “subject” variable was assigned as a random effect. Not all participants completed the scans for all three timepoints; however, we included participants who did not complete all three scan timepoints since a mixed effects modeling approach was applied, which is robust to missing data and accounts for the multiple timepoints of data (Yu et al., 2022). We fit different types of developmental trajectories to the breath-hold response, including linear, quadratic, logarithmic, and quadratic-log functions. The log, quadratic, and quadratic-log trajectories are common non-linear terms defined in developmental brain studies, as quadratic and logarithmic terms can allow for the characterization of peaks and plateaus in maturation times, respectively (Bethlehem et al., 2022; Faghiri et al., 2019). These trajectories have been previously reported in developmental studies (Herting et al., 2018). We implemented the ‘lme4’ package in R, and transformed the ‘age’ variable for quadratic, log, and quadratic-log functions (Bates et al., 2015). In addition to age, the duration of the breath-hold (BH) task was also added as a fixed effect, since some participants followed a 15s breath-hold task paradigm whereas others followed an 18s breath-hold task paradigm. Since the longitudinal cohort only has three timepoints, a random-intercept-only mixed effects model was used. Generally, a random slope and intercept model would require at least four timepoints to be reliable, particularly for higher-order models (King et al., 2018). The mixed effects models were formatted as follows:

Linear: CVR ∼ Age + BH Duration + (1|subject)

Quadratic: CVR ∼ Age^2^ + Age + BH Duration + (1|subject)

Logarithmic: CVR ∼ log(Age) + BH Duration + (1|subject)

Quadratic-log: CVR ∼ log(Age)^2^ + log(Age) + BH Duration + (1|subject)

### 2.7. Model Evaluation

We compared the different model types for each functional brain network. The linear, quadratic, log, and quadratic-log models were compared with each other to see which one provided the best fit, based on Akaike’s Information Criterion (AIC) score. The AIC score was calculated based on a maximum likelihood algorithm. We chose the AIC score as an evaluation of the goodness-of-fit of the models since no test-dataset was used and the models were all trained on the same data. The best-fitting model was chosen as the one with the lowest AIC score, which corresponds to a lower level of information loss (Portet, 2020).

### 2.8. Sex Effects

For each functional brain network, sex effects were evaluated by adding sex as a covariate in the mixed effects models for each brain network and each type of model: linear, quadratic, logarithmic, and quadratic logarithmic. The interaction between sex and age were also included in the mixed effects models to test for any interaction effects. T-tests using Satterthwaite’s method were used to evaluate the significance of the mixed effects model terms (‘lmerModLmerTest’ in R) (Kuznetsova et al., 2017).

## 3. Results

There was a total of 90 HC subjects who participated in the 18s BH paradigm during the BAS1 session, 82 HC subjects during the FLU1 session, and 60 HC subjects during the FLU2 session. For the 15s BH paradigm, there were 44 HC for BAS1, 12 HC for FLU1, and 1 HC subject for the FLU2 session. Not all subjects in the FLU1 groups completed a BAS1 MRI scan, therefore there were only a total of 49 HC subjects who completed both BAS1 and FLU1 sessions for the 18s BH paradigm and a total of 10 HC subjects who completed both BAS1 and FLU1 scans for the 15s BH paradigm. A total of 22 HC subjects completed all three sessions for the 18s BH paradigm and 0 subjects from the 15s BH paradigm group completed all three scans. Some subjects only had MRI data for one of the three timepoints (either BAS1, FLU1, or FLU2). After eliminating data due to head-motion and those with an insufficient number of timepoints, the data are as follows: 60 BAS1, 55 FLU1, and 40 FLU2 subjects for the 18s BH paradigm and 35 BAS1, 11 FLU1, and 0 FLU2 for the 15s BH paradigm (Figure 3). After head-motion correction, we further noticed that 12 subjects who performed the 15s BH paradigm for the BAS1 session had performed 18s BH for the FLU1 session, therefore we only kept the FLU1 scans for those participants, since it was the most recent scan, and we did not want to model the within-subject factors from both timepoints due to inconsistencies in the BH paradigm. This changed the number of subjects in the 15s BH paradigm during BAS1 from 35 to 23. Finally, all subject scans were visually inspected to check for any bad data quality or artifacts. During this stage, 6 subjects were excluded (from 15s BH: 1 BAS1 and 1 FLU1, from 18s BH: 1 BAS1, 1 FLU1, and 2 FLU2). Therefore, altogether, including both 15s and 18s BH paradigms, the total number of subject scans totaled 183 (Figure 3).

**Figure 3.**
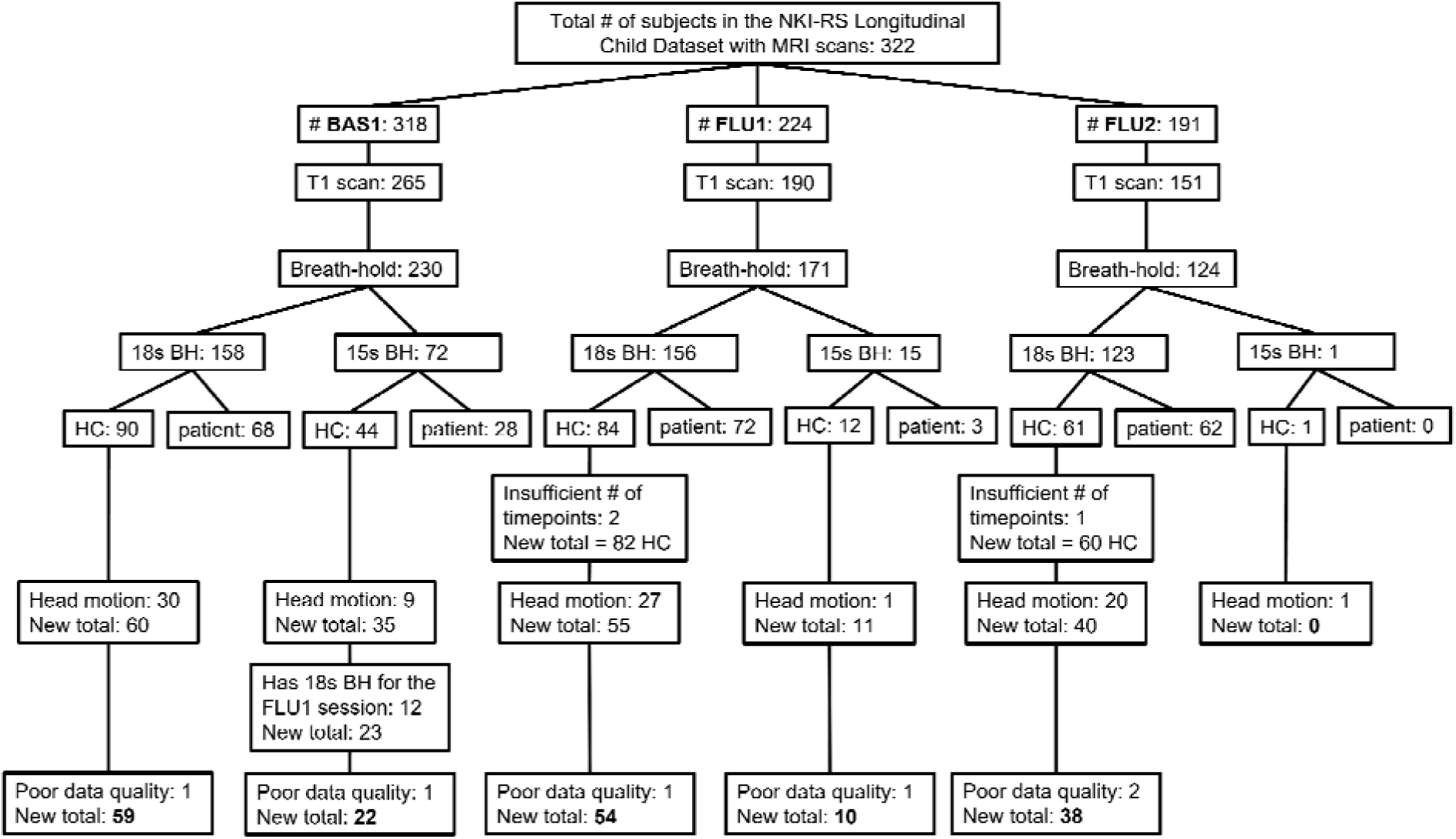
A flowchart showing the number of subjects in the NKI-RS Longitudinal Child Dataset with MRI scans for each session: baseline (BAS1), follow-up 1 (FLU1), and follow-up 2 (FLU2). Not all subjects started with MRI scans at the BAS1 session, some only have FLU1 or FLU2 scans. The final total number of subjects are highlighted in bold font.

Using ICA, the breath-hold data were decomposed into 12 identifiable functional brain networks for further analyses. All 12 IC networks are displayed using a “winner-takes-all” approach and the images were threshold at a z-score greater than 3. The following networks were defined: anterior and posterior default mode network, auditory, cingulo-opercular, frontal pole, lateral visual, left and right frontoparietal, medial visual, salience, sensorimotor, and somatosensory networks (Figure 4).

**Figure 4.**
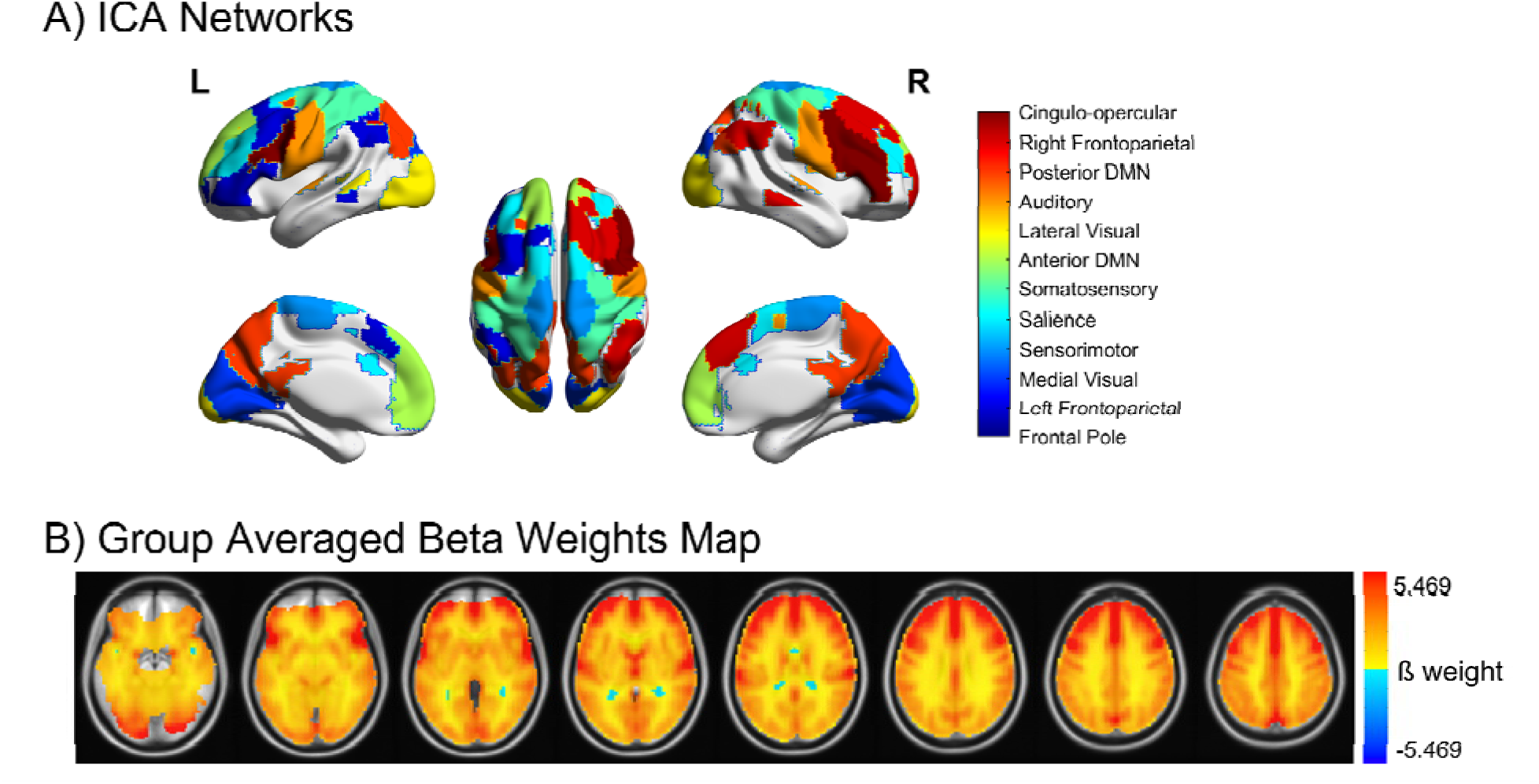
**A)** Map of all brain networks, visualized using BrainNetViewer (Xia et al., 2013). The IC maps are threshold at a z-score greater than 3 and displayed via a “winner-takes-all” approach. **B)** Group averaged beta weights map obtained using a general linear model (GLM) for all subjects.

### 3.1. CVR and Age

When accounting for all sessions of data (BAS1, FLU1, FLU2), and the BH duration, there were significant positive effects of age on CVR in the following networks: anterior DMN, auditory, cingulo-opercular, frontal pole, lateral visual, left frontoparietal, medial visual, and salience networks (Satterthwaite’s t-test, p<0.05 for all networks) (Figure 5 & Table 1). FDR correction was performed on the significant networks after choosing the model of “best-fit” for each network (see Section 3.2. Model Evaluation). Across all networks, the within-individual patterns visibly varied across time, showing that there may be low stability within individuals (Figure 5 (Crone and Elzinga, 2015). There were no significant effects of BH duration (18s vs 15s) on CVR across the 12 “best-fit” networks (Satterthwaite’s t-test, FDR-corrected p > 0.05) (Table S2).

**Figure 5.**
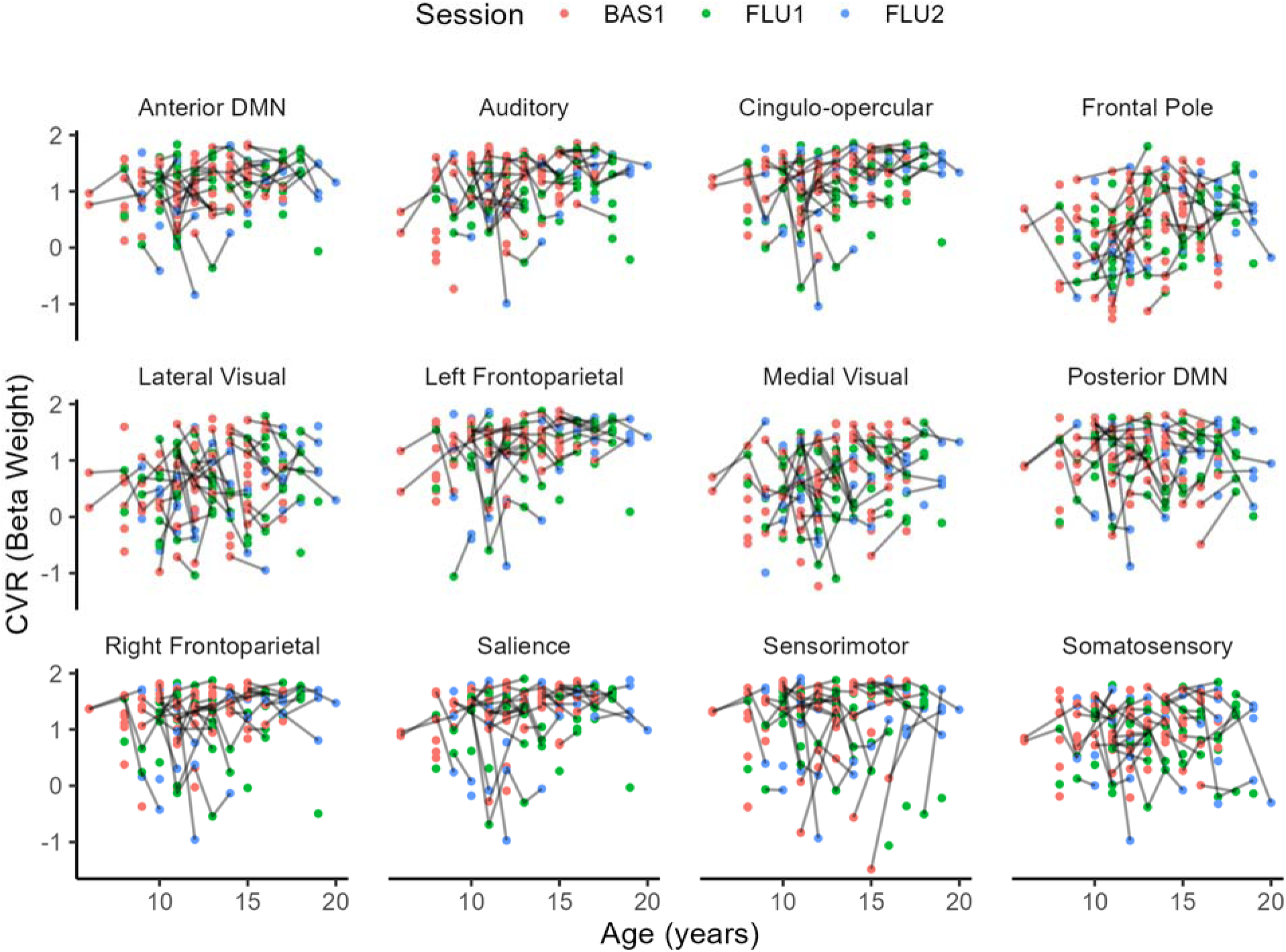
Relationship between cerebrovascular reactivity (CVR) and age for each brain network. Each gray line connecting the points represents a single subject’s data, connecting from one session to another. The age ranged from 6 to 20 years of age and not all subjects completed all three sessions. Some subjects only had one timepoint of scan, shown by a single point with no connecting lines. BAS1 = baseline (red), FLU1 = first follow-up (green), and FLU2 = second follow-up scan (blue).

**Table 1.** Mixed effects model evaluation for functional brain networks showing significant age effects. Estimates were obtained using the maximum likelihood algorithm. Model types highlighted in bold font indicate the best fitting model with the lowest AIC score for each network. FDR correction was performed on the corresponding p-values for each best-fit model.

| Brain Network | Model Type | Fixed Effect Estimate | 95% Confidence Interval | AIC Score | p-value for Age | p-value for BH |
| --- | --- | --- | --- | --- | --- | --- |

|  |  | (Age, Age<sup>2</sup>,<br>log(Age),<br>log(Age)<sup>2</sup>) | (Lower Bound,<br>Upper Bound) |  |  | duration |
| --- | --- | --- | --- | --- | --- | --- |
| Anterior DMN | Linear | 0.02742 | 0.00221,<br>0.05215 | 219.54<br>51 | 0.0288 | 0.129 |
|  | Quadratic | -0.00238 | -0.008809,<br>0.004051 | 221.01<br>41 | 0.467 | 0.301 |
|  | **Log** | 0.3481 | 0.03462, 0.6561 | 219.35<br>77 | 0.0259*<br>(FDR<br>adjusted:<br>0.0458) | 0.137 |
|  | Quadratic-Log | 0.02271 | -0.8495, 0.8965 | 221.35<br>51 | 0.959 | 0.915 |
| Auditory | Linear | 0.03432 | 0.006588,<br>0.06136 | 275.85<br>51 | 0.0128 | 0.00468 |
|  | **Quadratic** | -0.00836 | -0.01577, -<br>0.0009589 | 272.96<br>52 | 0.0267*<br>(FDR<br>adjusted:<br>0.0458) | 0.0115 |
|  | Log | 0.4809 | 0.139, 0.8159 | 274.21<br>93 | 0.00519 | 0.00537 |
|  | Quadratic-Log | -0.9205 | -1.94, 0.0978 | 273.07<br>49 | 0.0736 | 0.0491 |
| Cingulo-opercular | **Linear** | 0.04311 | 0.01641,<br>0.06933 | 266.24<br>52 | 0.00141*<br>(FDR<br>adjusted:<br>0.01002) | 0.0851 |
|  | Quadratic | 0.0002 | -0.007103,<br>0.00748 | 268.24<br>23 | 0.957 | 0.701 |
|  | Log | 0.5218 | 0.1879, 0.8504 | 266.82<br>86 | 0.00203 | 0.0903 |
|  | Quadratic-Log | 0.4927 | -0.5073, 1.49 | 267.88<br>77 | 0.329 | 0.444 |
| Frontal Pole | **Linear** | 0.05076 | 0.01572,<br>0.08545 | 348.35<br>87 | 0.0045*<br>(FDR<br>adjusted:<br>0.018) | 0.378 |
|  | Quadratic | -0.0011 | -0.0103,<br>0.008111 | 350.30<br>32 | 0.813 | 0.521 |
|  | Log | 0.6086 | 0.1708, 1.042 | 348.96<br>39 | 0.00635 | 0.368 |
|  | Quadratic-Log | 0.7588 | -0.4804, 2 | 349.51<br>45 | 0.229 | 0.314 |
| Lateral Visual | Linear | 0.03681 | 0.001802,<br>0.07105 | 359.06<br>64 | 0.0349 | 0.797 |
|  | Quadratic | -0.00304 | -0.01262,<br>0.006494 | 360.67<br>34 | 0.524 | 0.359 |
|  | **Log** | 0.4698 | 0.03529, 0.8969 | 358.82<br>81 | 0.0314*<br>(FDR<br>adjusted:<br>0.0471) | 0.778 |
|  | Quadratic-Log | -0.0885 | -1.401, 1.222 | 360.81<br>05 | 0.892 | 0.78 |
| Left<br>Frontoparietal | Linear | 0.04195 | 0.01445,<br>0.06893 | 268.80<br>24 | 0.00244 | 0.207 |
|  | Quadratic | -0.00411 | -0.01144,<br>0.003211 | 269.58<br>29 | 0.27 | 0.131 |
|  | **Log** | 0.5428 | 0.2007, 0.8789 | 268.11<br>22 | 0.00167*<br>(FDR<br>adjusted:<br>0.01002) | 0.223 |
|  | Quadratic-Log | -0.1887 | -1.19, 0.8128 | 269.97<br>44 | 0.711 | 0.559 |
| Medial Visual | Linear | 0.0394 | 0.006575,<br>0.07135 | 337.26<br>68 | 0.0155 | 0.0358 |
|  | Quadratic | -0.0031 | -0.0122,<br>0.005963 | 338.81<br>5 | 0.489 | 0.313 |
|  | **Log** | 0.4981 | 0.09144, 0.8965 | 337.04<br>07 | 0.0146*<br>(FDR<br>adjusted:<br>0.03504) | 0.0377 |
|  | Quadratic-Log | -0.01655 | -1.278, 1.244 | 339.04 | 0.978 | 0.849 |
| Salience | Linear | 0.03507 | 0.007629,<br>0.06198 | 266.07<br>17 | 0.0107 | 0.279 |
|  | Quadratic | -0.00335 | -0.01064,<br>0.003932 | 267.25<br>12 | 0.366 | 0.212 |
|  | **Log** | 0.4493 | 0.1075, 0.785 | 265.72<br>48 | 0.0088*<br>(FDR<br>adjusted:<br>0.0264) | 0.294 |
|  | Quadratic-Log | -0.04512 | -1.04, 0.9498 | 267.71<br>68 | 0.929 | 0.789 |

### 3.2. Model Evaluation

Based on Akaike’s Information Criterion (AIC) scores, different age-trajectories in the mixed effects models provided better fits for different brain networks (Table 1 & Figure S1). Focusing on the eight brain networks with significant age effects, we compared the different age-trajectory models to see which fit best. For these significant networks, the best-fitting model was chosen as the one with the lowest AIC score (Burnham and Anderson, 2004). Out of the networks which had significant age effects on CVR after FDR correction, the logarithmic model appeared most frequently as the “best-fit” model, for the following networks: anterior DMN, lateral visual, left frontoparietal, medial visual, and salience networks. The linear age trajectory model provided the best fit for the cingulo-opercular and frontal pole networks, and the quadratic model provided the best fit for the auditory network. The quadratic-log model did not appear as the “best-fit” model for any of the brain networks (Table 1).

### 3.3. Sex differences

We found no significant effects of sex on CVR for each mixed effects model, and no interaction effects between sex and age. For all brain networks, there were no significant effects of sex found. Across the 12 networks, the trajectories between males and females were similar and overlapped (Figure 6).

**Figure 6.**
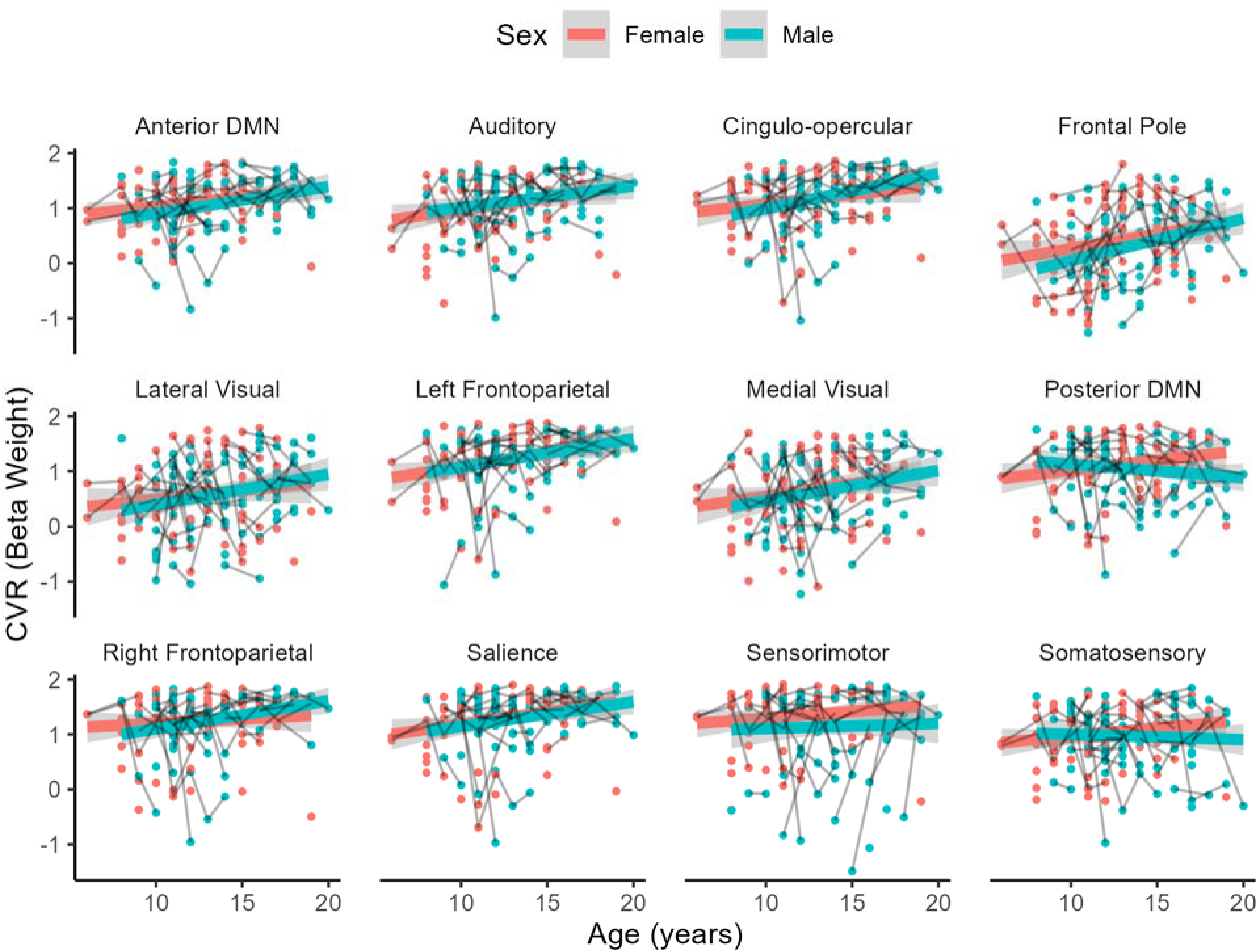
CVR (beta weight) values plotted across age for all 12 networks, with linear trajectories for females and males shown separately. The dark gray lines connecting the points represent a single subject at different sessions.

## 4. Discussion

In the current study, we fit linear, logarithmic, quadratic, and quadratic-logarithmic mixed effects models to longitudinal breath-hold response data across age, from 6 to 20 years. We found that different functional brain networks exhibited different breath-hold response trajectories across age, with linear and logarithmic models providing the best fit. In particular, the anterior DMN, lateral visual, left frontoparietal, medial visual, and salience networks displayed significant logarithmic relationships in CVR across age, while the cingulo-opercular and frontal pole networks showed significant linear age-trajectory relationships, and the auditory network with significant quadratic age-trajectory in CVR. We report no sex-effects on the breath-hold response across age.

### 4.1. Increased cerebrovascular reactivity across multiple networks

Different functional brain networks have been shown to mature at different rates across age (Gaillard et al., 2001; Zuo et al., 2010). We observed an increase in cerebrovascular reactivity in multiple networks across age, with varying linear, quadratic, and logarithmic trajectories. The breath-hold response is a global response, so it influences all regions of the brain. However, it appears that across development, there are certain areas of the brain that are more susceptible to the breath-hold response or have different patterns of response.

The medial visual and lateral visual networks both showed significant logarithmic trends in CVR across age. Marcar and colleagues found that the BOLD response in younger children is more strongly influenced by cerebral metabolic rate of oxygen (CMRO2) and suggest that the visual system in young children is immature, thus less energy efficient than a mature visual system (Marcar et al., 2004). As age increases, CVR was seen to increase as well, possibly indicative of the development of a more mature visual system. In animal studies, visual enrichment has been shown to increase the density of vessels per neuron (Argandoña et al., 2012). As children age, the visual system is exposed to more stimuli and thus the vasculature in the visual networks can be expected to increase. Increased vascular density in these areas may be the cause for increased CVR across age. Since the breath-hold task causes vasodilation, a denser vascular architecture may lead to an increased breath-hold response. Increased vascular maturation in the visual cortex would also lead to increased local cerebral blood flow (CBF) and cerebral blood volume (CBV), which both contribute to cerebrovascular reactivity (Bouma and Muizelaar, 1992).

Similarly, the frontal pole network showed increased CVR across age, which supports our hypothesis that CVR would increase in the frontal region until late adolescence. The frontal pole network is associated with higher-order cognitive areas and tends to mature later in life, particularly since this region is mediated by experience. Urback and colleagues found significant regional differences in CVR in healthy adolescents, with the highest CVR found in the frontal and occipital regions, followed by the parietal, temporal, and then other MNI regions (Urback et al., 2018). The primary sensory and motor areas of the brain complete development first, whereas the association areas are the last to mature (Gaillard et al., 2001). In a longitudinal fMRI study, Ordaz and colleagues reported that motor-related regions showed no age-related changes, while error-related activity increased into early adulthood in the anterior cingulate cortex (Ordaz et al., 2013; Telzer et al., 2018). This is consistent with our findings, in which the motor-related networks showed no significant age-related changes in cerebrovascular reactivity. Many of the networks which showed significant effects of age on CVR were part of the association cortex, such as the frontal pole, frontoparietal, auditory, medial visual, and lateral visual networks, which receive inputs from multiple areas of the brain. This suggests a possible hierarchical structure in CVR development, with higher-order regions continuing to develop later in life.

We found a significant logarithmic age-effect on CVR in the anterior DMN region, but no significant age-effects in the posterior DMN region. The anterior DMN is involved in conscious planning and control functions whereas the posterior DMN is more involved in unconscious self-representation and self-centered cognitive functions (Knyazev, 2012). The distinction in age-effects between these two DMN regions indicate that CVR developmental trajectories are unique and specific. However, CVR may also not be related to function since breath-hold fMRI is different from typical task-based fMRI in that it is primarily a vascular stimulus. Therefore, associating the breath-hold response in different brain regions to the function of those particular brain regions may not be applicable. It is not yet clear why specific networks showed age-effects in CVR; however, it is possible that this is dependent on the cerebrovascular structure and some areas may have a more dense vascular network than others. We found that CVR is altered across age in brain network-specific patterns, similar to how brain development has been shown to be network-specific.

Across all networks, we observed state-like variability within individuals. Cross-sectional fMRI studies on development may not be reflective of development, particularly when there are intra-individual variabilities that can only be studied with longitudinal data (Crone and Elzinga, 2015). The data used in the present study is longitudinal in design; however, there are only 3 separate scan sessions and not all subjects completed all scan sessions due to the challenge of subject attrition. Many fMRI studies investigating the developing brain are cross-sectional in design and this limits our understanding of development, as each child’s brain is unique and may follow different trajectories of development. A cross-sectional design may overlook these unique developmental trajectories. Despite the advantages of longitudinal developmental study designs, there are challenges which limit its wide-spread usage. It is not only difficult to have participants commit to follow-up scans, but these study designs may also be very costly. Another challenge arises during the consideration of a specific task to study developmental cognitive functions.

When a participant is presented with a cognitive or behavioral task, they typically respond the strongest during the initial exposure to the task. As the task is repeated multiple times, the participant may become accustomed to the task or have expectations ready, which may yield weaker responses. This is important to consider when designing longitudinal studies, since any effects observed could simply be due to habituation. However, this does not apply to the breath-hold task since it induces hypercapnia and there are minimal habituation effects associated with hypercapnia. In addition to task-based fMRI studies, resting-state studies are also widely used in longitudinal designs, in which the participant is not asked to perform any specific task, but rather lay in the fMRI scanner in an awake and rested state. This type of design measures brain activity more intrinsically without any external stimuli, and in a longitudinal resting-state design, researchers do not have to worry about any habituation or learning effects. Yet despite the advantages of resting-state fMRI longitudinal studies, limitations also arise in terms of interpretability of the ongoing neuronal vs. vascular contributions to the blood-oxygen-level-dependent (BOLD) signal, which is yet unclear.

### 4.2. Trends that best describe cerebrovascular reactivity across children

We observed linear, quadratic, and logarithmic trajectories of CVR across age in the majority of networks investigated, such as the anterior DMN, auditory, cingulo-opercular, frontal pole, lateral visual, left frontoparietal, medial visual, and salience networks. Although there are no longitudinal breath-hold fMRI studies that we are aware of, there have been studies investigating longitudinal developmental trajectories using cognitive task-based fMRI across children. Ordaz and colleagues longitudinally investigated the performance of an inhibitory anti-saccade task in participants aged 9 to 26 years old and observed linear effects of age in activation in the supplementary eye fields, left posterior parietal cortex, and putamen areas (Ordaz et al., 2013).

Braams and colleagues investigated longitudinal changes across development in the nucleus accumbens, a region of the brain associated with rewards and risk taking and found a quadratic age pattern where a peak response to rewards was found in adolescence (Braams et al., 2015). These studies fitted linear and quadratic models to longitudinal fMRI data and chose the model which provided the best fit to describe development. Across the different brain networks, different models to describe the developmental trajectory were fitted, which shows that CVR across age may also develop differentially in different brain networks and tasks. These studies used task-based fMRI and did not include breath-hold paradigms; however, the breath-hold response has been suggested to contribute to the fMRI BOLD signal activity. Therefore, the trajectories stated in previous papers could also be due in part to underlying differences in CVR. The nonlinear positive trajectories observed in our study may also reflect puberty-dependent changes which occur in the brain. Gracia-Tabuenca and colleagues show that there are nonlinear puberty-dependent trajectories in functional brain organization across adolescents, particularly in the frontal and parietal functional networks (Gracia-Tabuenca et al., 2021). In particular, the onset of puberty is associated with increased synaptic pruning which occurs in the frontal areas (Drzewiecki et al., 2016; Huttenlocher, 1979).

Using a breath-hold paradigm, Thomason and colleagues reported significant differences between children (7-12 yrs) and adults (18-29 yrs) in the number of voxels activated across brain regions and between brain regions (Thomason et al., 2005). This showed heterogeneity in the vascular responsiveness across the brain, which was consistent with our findings of increased CVR in specific networks. Regional differences in the responsiveness of the brain to the breath-hold stimulus has been shown previously by Kastrup and colleagues (Kastrup et al., 1999a); however, Thomason and colleagues were the first to test the regional effect of breath-holding between two different age groups – children and adults. In the present study, we observed network-specific age trajectories of the breath-hold response in children enrolled in a longitudinal study.

### 4.3. No sex effects on cerebrovascular reactivity across children

For each brain network, no significant effect of sex on cerebrovascular reactivity was found. In Ordaz and colleagues’ longitudinal fMRI study of participants aged 9 to 26 years, there were sex-effects predominantly in the motor areas of the brain as well as the executive control region in the right ventrolateral prefrontal cortex (Ordaz et al., 2013). In contrast, we found no effect of sex on the breath-hold response in the motor and executive function areas of the brain for each model fit after FDR correction. In a diffusion tensor imaging (DTI) study investigating structural connectivity differences between males and females across development, females were observed to have interhemispheric connectivity dominance mainly in the frontal lobe during adolescence (defined as ages 13.4-17 years), whereas during adulthood (17.1-22 years), the interhemispheric connectivity dispersed to other lobes (Ingalhalikar et al., 2014).

### 4.4. Limitations

One limitation of the study is the lack of subjects for the younger age range. The younger aged group (<8 years) typically has higher levels of head motion. Since we don’t have measures of vascular activity for those younger than 6 years of age, it is possible that there was a logarithmic trend in many networks, but due to stabilization already occurring at 6 years of age, we cannot observe those logarithmic effects. Future studies may want to focus on more high-quality data in the younger-aged group to be more representative of development. Measures of subject compliance were also not provided for all subjects, and the measurements were very noisy, so it is not completely clear if the subjects performed the breath-hold task correctly.

Younger subjects may be less compliant with the breath-hold task; however, even poor performance on the breath-hold task has been shown to yield robust measurements of CVR (Bright and Murphy, 2013). Another limitation to consider when implementing the breath-hold challenge is the metabolic rate difference between participants and therefore different rate of oxygen consumption and carbon dioxide production, which makes the results less comparable between individuals. In children’s developmental studies, this is a potential confounder, particularly due to the changes in metabolic rate as a child grows and fat-free body mass increases (Pontzer et al., 2021). The NKI-RS longitudinal children study only provides BMI scores for one baseline timepoint, therefore we decided not to include this in our model; however, future studies may want to further explore the effect of BMI on CVR measures in children.

## 5. Conclusion

The current study investigated the longitudinal brain trajectories of development in children performing a breath-hold task, using fMRI. We observed that cerebrovascular reactivity across development follows a logarithmic trajectory overall; however, different functional brain networks may be explained better by different mixed models. We also report no sex-effects in the breath-hold response across brain networks and models.

## CRediT authorship contribution statement

**Donna Y. Chen:** conceptualization, formal analysis, writing. **Xin Di:** conceptualization, supervision, writing - review&editing. **Bharat Biswal:** conceptualization, supervision, funding acquisition, writing - review&editing.

## Acknowledgements

Research reported in this publication was supported by the National Institutes of Health (NIH) under Award Number R01MH012577 and RF1NS124778 to B.B., and R15MH125332 to X.D, and by the National Center for Advancing Translational Sciences (NCATS) of the NIH under Award Number TL1TR003019 to D.Y.C. The content is solely the responsibility of the authors and does not necessarily represent the official views of the National Institutes of Health.

