## SupplementaryFigures for "Cerebrovascular reactivity increases across development in multiple networks as revealed by a breath-holding task: a longitudinal fMRI study"

**Supplementary Tables/Figures**

**Table S1.** Mixed effects model evaluation for all brain networks. Estimates were obtained using the maximum likelihood algorithm. For each network, the model type that provides the best fit based on the AIC score is in bold font. For the best-fit models, a * indicates significance after BH-FDR correction.

| **Brain Network** | **Model Type** | **Fixed Effect Estimate (Age, Age^2^, log(Age), log(Age)^2^)** | **95% Confidence Interval (Lower Bound, Upper Bound)** | **AIC Score** | **p-value for Age** | **p-value for BH duration** |
| --- | --- | --- | --- | --- | --- | --- |
| Anterior DMN | Linear | 0.02742 | 0.00221, 0.05215 | 219.5451 | 0.0288 | 0.129 |
|  | Quadratic | -0.00238 | -0.008809, 0.004051 | 221.0141 | 0.467 | 0.301 |
|  | **Log** | 0.3481 | 0.03462, 0.6561 | 219.3577 | 0.0259 | 0.137 |
|  | Quadratic-Log | 0.02271 | -0.8495, 0.8965 | 221.3551 | 0.959 | 0.915 |
| Auditory | Linear | 0.03432 | 0.006588, 0.06136 | 275.8551 | 0.0128 | 0.00468 |
|  | **Quadratic** | -0.00836 | -0.01577, -0.0009589 | 272.9652 | 0.0267 | 0.0115 |
|  | Log | 0.4809 | 0.139, 0.8159 | 274.2193 | 0.00519 | 0.00537 |
|  | Quadratic-Log | -0.9205 | -1.94, 0.0978 | 273.0749 | 0.0736 | 0.0491 |
| Cingulo-opercular | **Linear** | 0.04311 | 0.01641, 0.06933 | 266.2452 | 0.00141 | 0.0851 |
|  | Quadratic | 0.0002 | -0.007103, 0.00748 | 268.2423 | 0.957 | 0.701 |
|  | Log | 0.5218 | 0.1879, 0.8504 | 266.8286 | 0.00203 | 0.0903 |
|  | Quadratic-Log | 0.4927 | -0.5073, 1.49 | 267.8877 | 0.329 | 0.444 |
| Frontal Pole | **Linear** | 0.05076 | 0.01572, 0.08545 | 348.3587 | 0.0045 | 0.378 |
|  | Quadratic | -0.0011 | -0.0103, 0.008111 | 350.3032 | 0.813 | 0.521 |
|  | Log | 0.6086 | 0.1708, 1.042 | 348.9639 | 0.00635 | 0.368 |
|  | Quadratic-Log | 0.7588 | -0.4804, 2 | 349.5145 | 0.229 | 0.314 |
| Lateral Visual | Linear | 0.03681 | 0.001802, 0.07105 | 359.0664 | 0.0349 | 0.797 |
|  | Quadratic | -0.00304 | -0.01262, 0.006494 | 360.6734 | 0.524 | 0.359 |
|  | **Log** | 0.4698 | 0.03529, 0.8969 | 358.8281 | 0.0314 | 0.778 |
|  | Quadratic-Log | -0.0885 | -1.401, 1.222 | 360.8105 | 0.892 | 0.78 |
| Left Frontoparietal | Linear | 0.04195 | 0.01445, 0.06893 | 268.8024 | 0.00244 | 0.207 |
|  | Quadratic | -0.00411 | -0.01144, 0.003211 | 269.5829 | 0.27 | 0.131 |
|  | **Log** | 0.5428 | 0.2007, 0.8789 | 268.1122 | 0.00167 | 0.223 |
|  | Quadratic-Log | -0.1887 | -1.19, 0.8128 | 269.9744 | 0.711 | 0.559 |
| Medial Visual | Linear | 0.0394 | 0.006575, 0.07135 | 337.2668 | 0.0155 | 0.0358 |
|  | Quadratic | -0.0031 | -0.0122, 0.005963 | 338.815 | 0.489 | 0.313 |
|  | **Log** | 0.4981 | 0.09144, 0.8965 | 337.0407 | 0.0146 | 0.0377 |
|  | Quadratic-Log | -0.01655 | -1.278, 1.244 | 339.04 | 0.978 | 0.849 |
| Posterior DMN | Linear | -0.00186 | -0.02977, 0.02586 | 288.544 | 0.895 | 0.316 |
|  | **Quadratic** | -0.00575 | -0.01343, 0.00192 | 288.3758 | 0.141 | 0.15 |
|  | Log | 0.01561 | -0.3325, 0.3619 | 288.5535 | 0.929 | 0.327 |
|  | Quadratic-Log | -0.7451 | -1.796, 0.3054 | 288.6112 | 0.164 | 0.163 |
| Right Frontoparietal | **Linear** | 0.02726 | -0.001093, 0.05501 | 283.4604 | 0.0532 | 0.126 |
|  | Quadratic | 0.000311 | -0.007323, 0.007936 | 285.454 | 0.936 | 0.854 |
|  | Log | 0.3235 | -0.03124, 0.671 | 283.8102 | 0.0666 | 0.129 |
|  | Quadratic-Log | 0.4812 | -0.5617, 1.523 | 284.9852 | 0.364 | 0.433 |
| Salience | Linear | 0.03507 | 0.007629, 0.06198 | 266.0717 | 0.0107 | 0.279 |
|  | Quadratic | -0.00335 | -0.01064, 0.003932 | 267.2512 | 0.366 | 0.212 |
|  | **Log** | 0.4493 | 0.1075, 0.785 | 265.7248 | 0.0088 | 0.294 |
|  | Quadratic-Log | -0.04512 | -1.04, 0.9498 | 267.7168 | 0.929 | 0.789 |
| Sensorimotor | Linear | 0.002482 | -0.03049, 0.03539 | 364.7256 | 0.882 | 0.0135 |
|  | Quadratic | -0.00452 | -0.01383, 0.004826 | 365.8177 | 0.34 | 0.335 |
|  | **Log** | 0.05855 | -0.3538, 0.47 | 364.669 | 0.779 | 0.014 |
|  | Quadratic-Log | -0.5194 | -1.8, 0.7635 | 366.0323 | 0.425 | 0.416 |
| Somatosensory | Linear | -0.00068 | -0.02961, 0.02781 | 292.7365 | 0.962 | 0.158 |
|  | **Quadratic** | -0.0061 | -0.01396, 0.00174 | 292.4033 | 0.125 | 0.131 |
|  | Log | 0.02637 | -0.333, 0.3815 | 292.7175 | 0.884 | 0.165 |
|  | Quadratic-Log | -0.5987 | -1.672, 0.4731 | 293.5108 | 0.27 | 0.267 |

**Table S2.** Models are chosen based on the best-fit (lowest AIC score). The p-values are adjusted using Benjamini-Hochberg’s FDR correction. There are 8 significant networks after FDR correction, of which 2 are linear, 1 is quadratic, and 5 are logarithmic.

| **IC network** | **Best-fit model** | **FDR-adjusted p-val** | |
| --- | --- | --- | --- |
|  |  | **Age** | **BH duration** |
| Anterior DMN | Log | 0.045771* | 0.225 |
| Auditory | Quadratic | 0.045771* | 0.084 |
| Cingulo-opercular | Linear | 0.01002* | 0.225 |
| Frontal Pole | Linear | 0.018* | 0.412364 |
| Lateral Visual | Log | 0.0471* | 0.778 |
| Left Frontoparietal | Log | 0.01002* | 0.297333 |
| Medial Visual | Log | 0.03504* | 0.1508 |
| Posterior DMN | Quadratic | 0.153818 | 0.225 |
| Right Frontoparietal | Linear | 0.070933 | 0.225 |
| Salience | Log | 0.0264* | 0.3528 |
| Sensorimotor | Log | 0.779 | 0.084 |
| Somatosensory | Quadratic | 0.15 | 0.225 |

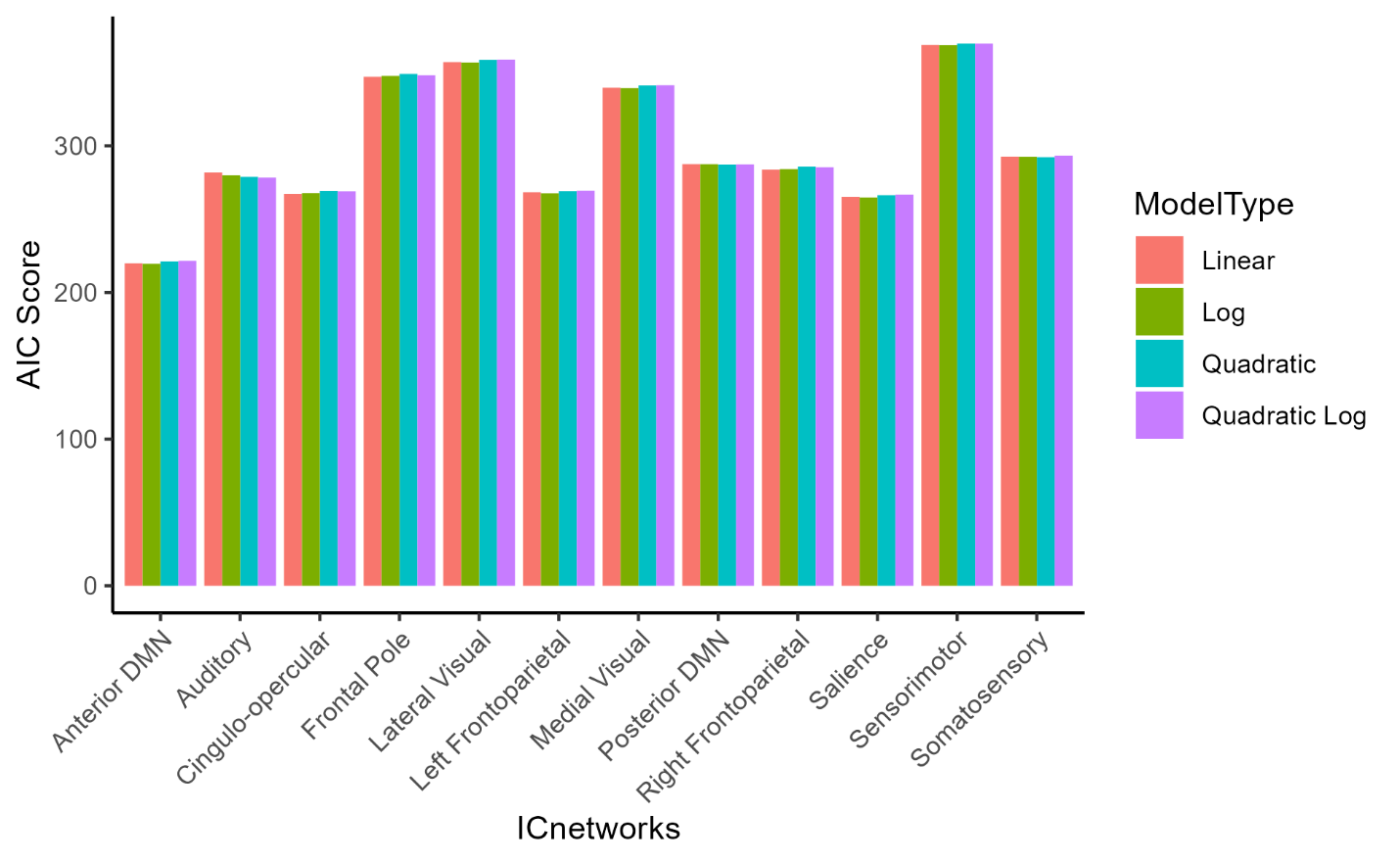

**Figure S1.** Mixed effects model evaluation for each brain network. Estimates were obtained using the maximum likelihood algorithm. The AIC scores for each IC brain network, and each model type: linear, log, quadratic, and quadratic-log are shown. A lower AIC score represents a better goodness of fit for the model.
